# Stronger Ca²⁺-to-voltage coupling in females promotes cellular alternans but limits spatial discordance by favoring electrotonic phase-locking

**DOI:** 10.64898/2026.11.19.767741

**Authors:** Roshni Shetty, Lianguo Wang, Daisuke Sato, Jason D. Bayer, Crystal M. Ripplinger, Eleonora Grandi

## Abstract

Repolarization alternans, a beat-to-beat oscillation in action potential duration (APD), reflects electrical instability associated with ventricular arrhythmias and sudden cardiac death. Often driven by Ca²⁺ transient (CaT) alternans, its magnitude also depends on Ca²⁺-to-voltage coupling through Ca²⁺-sensitive membrane currents. β-Adrenergic signaling (βARS) modulates repolarization, Ca²⁺ handling and alternans, and becomes spatially heterogeneous in diseased hearts. Sex differences in repolarization and Ca²⁺ handling further influence arrhythmia susceptibility. How these factors interact to shape spatial alternans remains unclear.

We simulated sex-specific human ventricular myocyte models incorporating βARS in isolated cells and one-dimensional strands containing a local βARS-unstimulated region. We examined how sex, βARS, Ca²⁺-to-voltage coupling, and electrotonic coupling influence alternans, and compared predictions with rabbit ventricular optical mapping data.

Alternans required beat-to-beat alternation in SR Ca²⁺ loading and was translated into APD alternans primarily through Na⁺/Ca²⁺ exchanger (INaCa). Female myocytes exhibited larger APD alternans despite smaller CaT alternans, reflecting stronger Ca²⁺-to-voltage coupling. In coupled strands, however, this higher coupling gain enhanced electrotonic phase-locking, preserving APD concordance despite local CaT discordance. Reducing female INaCa to male levels, weakening electrotonic coupling, or enlarging unstimulated regions abolished this protection. Optical mapping showed a qualitatively similar dissociation between CaT discordance and preserved APD concordance.

We conclude that Ca²⁺-to-voltage coupling can amplify cellular APD alternans but suppress spatial discordance. This effect depends on the spatial extent of heterogeneity and may be diminished by fibrosis, gap-junction remodeling, or heterogeneous innervation. Thus, arrhythmic risk may depend not only on sympathetic drive but also on its spatial distribution.

**NEW & NOTEWORTHY:** Using sex-specific human ventricular action potential models incorporating βARS, we show that female vs. male myocytes develop larger repolarization alternans despite smaller Ca²⁺ transient alternans, because higher INaCa-dependent Ca²⁺-to-voltage gain amplifies APD alternans. In one-dimensional strands, the same gain limits APD discordance: large APD alternans in flanking stimulated regions electrotonically entrains a local βARS unstimulated region, preserving APD concordance despite Ca²⁺ discordance. Weaker electrotonic coupling or a larger unstimulated region abolish this protection.

## 1. INTRODUCTION

Sudden cardiac death (SCD) remains a leading cause of mortality worldwide and most commonly results from ventricular tachycardia (VT) that degenerates into ventricular fibrillation (VF) (1–3). Repolarization alternans, defined as beat-to-beat alternation of action potential duration (APD), is an established marker of electrical instability that manifests on the ECG as T-wave alternans (TWA) and predicts ventricular arrhythmias and SCD (2, 4–6).

At the cellular level, APD alternans often originates from instability of intracellular Ca²⁺ cycling. At rapid rates, beat-to-beat differences in sarcoplasmic reticulum (SR) Ca²⁺ load and release, incomplete recovery of ryanodine receptors (RyR2), or interactions between these mechanisms can produce Ca²⁺ transient (CaT) alternans (7–11). CaT alternans is coupled to membrane voltage through Ca²⁺ sensitive currents. In particular, the Na⁺/Ca²⁺ exchanger (INaCa) converts beat-to-beat differences in Ca²⁺ extrusion into changes in membrane current, while the L-type Ca²⁺ current and Ca²⁺-sensitive K⁺ currents further set the sign and magnitude of this coupling (12–15). Consequently, the magnitude of APD alternans depends both on the degree of instability in Ca²⁺ cycling and on the gain with which a given ΔCaT is translated into ΔAPD. Consistent with this framework, impaired SR Ca²⁺ uptake, including reduced SERCA activity in heart failure, increases alternans susceptibility by shifting its onset toward slower pacing rates (16–18).

Critical to arrhythmogenesis is the spatial organization of alternans. When neighboring regions alternate out of phase, spatially discordant alternans (SDA) create steep repolarization gradients that promote unidirectional block, wavebreak and reentrant VF (2, 8, 19, 20). SDA can emerge dynamically from steep APD and conduction velocity (CV) restitution (19), but it can also develop around pre-existing tissue heterogeneity, including electrotonic uncoupling at a structural barrier (20, 21) or impaired gap-junction coupling (22–24). Critically, voltage and Ca²⁺ organize over different spatial scales. Ca²⁺ alternans can reverse phase over roughly a single cell length, whereas electrotonic coupling smooths APD over millimeters (25–27). Accordingly, CaT and APD nodal lines need not coincide (28). Whether a local Ca²⁺ phase reversal manifests as APD discordance therefore depends on the balance between the local Ca²⁺ instability and entrainment by the surrounding myocardium, as well as on Ca²⁺-to-voltage gain and the spatial extent of the heterogeneous region (29).

β-adrenergic receptor signaling (βARS) provides a physiologically important setting in which these mechanisms may interact (30). In isolated myocytes and intact hearts, β-AR stimulation typically suppresses Ca²⁺ and repolarization alternans, largely by accelerating SR Ca²⁺ reuptake via phospholamban phosphorylation and by modulating SR Ca²⁺ release (10, 31–33). In diseased hearts, however, sympathetic innervation becomes markedly non-uniform, with nerve sprouting adjacent to denervated regions (34), and the severity of these innervation gradients correlates with arrhythmia (Dajani *et al.*, 2023). Regional differences in sympathetic input alter local repolarization and increase repolarization dispersion (35), and simulations show that increasing spatial heterogeneity in βARS is associated with greater arrhythmogenicity (36, 37). Thus, a signaling pathway that suppresses alternans locally may nevertheless generate spatial differences in alternans susceptibility when its activation is nonuniform.

Sex may further modify this relationship. Females have lower rates of SCD and sustained ventricular arrhythmias than males (38, 39), yet exhibit longer QT intervals and greater susceptibility to drug-induced torsades de pointes (40, 41). Sex differences in ion channel expression and Ca²⁺ handling may alter both the development of cellular alternans and the coupling of Ca²⁺ alternans to repolarization. Whether these cellular differences translate into corresponding sex differences in the spatial organization of alternans, particularly in the presence of heterogeneous βARS, remains unclear.

Here, we combined sex-specific human ventricular myocyte models incorporating βARS with one-dimensional multicellular strands containing a localized region without βARS, and compared key model predictions with simultaneous voltage and Ca²⁺ optical mapping in rabbit ventricle. We sought to determine: (i) how sex affects cellular alternans susceptibility and its modulation by βARS and SR Ca²⁺ uptake, and (ii) how spatial heterogeneity in βARS governs the emergence and spatial organization of alternans in tissue. We hypothesized that spatial APD patterns reflect a competition between locally driven Ca²⁺ phase reversal and electrotonic phase-locking by the surrounding myocardium, and that sex-dependent differences in Ca²⁺-to-voltage coupling alter the balance between these opposing influences.

## 2. MATERIALS AND METHODS

### 2.1 Sex-specific human ventricular myocyte model with β-adrenergic signaling

Cellular electrophysiology was simulated with the O’Hara-Rudy (ORd) model of the human ventricular endocardial myocyte (42). The maximal conductances of IKr, IKs, IK1, ICaL and INaL were scaled following the CiPA optimization by (43). The ORd fast Na⁺ current was replaced by the ten Tusscher-based (44) INa formulation used in the (45) model to support excitability and conduction in multicellular simulations. Male and female phenotypes were implemented as in (46), based on human sex differences in ion-channel expression and previous models (40, 47, 48). β-adrenergic signaling was incorporated using the (49) implementation of the compartmentalized (50) signaling network coupled to the ORd model, with INa phosphorylation adapted from (51). The network describes isoproterenol (ISO) binding to β₁- and β₂-adrenergic receptors, adenylyl cyclase activation, cAMP production and degradation by phosphodiesterases (PDE2-4), protein kinase A (PKA) activation in caveolar, extracaveolar and cytosolic compartments. PKA phosphorylates eight targets: caveolar: L-type Ca²⁺ channel, fast Na⁺ channel, Na⁺/K⁺ pump, ryanodine receptor; extracaveolar: IKs and the background K⁺ current; cytosolic: phospholamban (PLB) and troponin I (TnI). Interactions with Ca^2+^-calmodulin, Protein kinase II CaMKII phosphorylation and protein phosphatase 1 PP1-mediated dephosphorylation were retained as in the original signaling model. No sex differences were implemented in the signaling pathway. To reduce computational cost, the signaling and electrophysiology modules were simulated in a decoupled manner. For each ISO concentration, the signaling model was first run to steady state to determine phosphorylation fractions for the eight PKA targets and whole-cell PP1 activity. These steady-state values were then fixed during the electrophysiological simulations.

### 2.2 Single-cell pacing protocols

Cells were stimulated with -80 µA µF⁻¹ current pulses of 0.5 ms. After 3 s of quiescence from the initial conditions (default (42) endocardial initial conditions), a dynamic pacing protocol was applied: 100 beats at each cycle length (CL); CL decreased from 600 ms to 500 ms in 50ms steps, then in 10 ms steps to 450 ms, and in 5 ms steps to 200 ms, or until loss of 1:1 capture. The last two beats at each CL (beats 99 and 100) were analyzed.

In some simulations, selected variables were clamped at CL = 245 ms to isolate specific mechanisms. For INaCa clamp and SR Ca²⁺ clamp, the model was first paced for 100 beats and the longer of beats 99 and 100 was identified. The corresponding INaCa,i and INaCa,ss waveforms, or [Ca²⁺]NSR and [Ca²⁺]JSR trajectories, were then imposed at every time step during a subsequent 100-beat simulation. For cross-sex Ca²⁺ clamps, the male [Ca²⁺]_i_ trajectory over 100 beats was imposed on the female model using linear interpolation; a male self-clamp served as control.

### 2.3 One-dimensional strand simulations

Male and female endocardial strands consisted of 400 cells (Δx = 125 µm; total length 5 cm). Propagation was described by the monodomain equation ∂V/∂t = D ∂²V/∂x² − Iion/Cm, with sealed (no-flux) boundaries. The diffusion coefficient D was 0.0010 cm² ms⁻¹ (normal coupling) or 0.0003 cm² ms⁻¹ (weak coupling), giving a conduction velocity of 65 and 33 cm s⁻¹ respectively at CL 600 ms.

With baseline SERCA, action potential propagation along the strand did not exhibit alternans. SERCA uptake was therefore reduced by 40% (×0.6) in all cells to represent a heart-failure-like Ca²⁺-handling phenotype (16, 18). To mimic a locally denervated region within sympathetically stimulated tissue, cells were assigned the phosphorylation fractions for 10 nM ISO, except cells 200-224 (2.50-2.81 cm), which received the 0 nM ISO fractions. Thus, the strand contained a central unstimulated region (ISO = 0 nM) embedded within flanking ISO-stimulated tissue (0-2.5 cm and 2.81-5 cm, ISO= 10 nM). All cells started from the same initial state (default (42) endocardial initial conditions) and rested for 3 s. The first two cells were then stimulated (-150 µA µF⁻¹, 1 ms) with a dynamic protocol of 100 beats per CL: CL decreased from 600 ms to 500 ms in 50 ms steps, then to 450 ms in 10 ms steps, and in 5 ms steps to 200 ms, or until loss of 1:1 capture, identical to the single cell.

### 2.4 Biomarkers and alternans definitions

#### APD and peak CaT

APD95 was measured from activation (maximal dV/dt) to 95% repolarization, and CaTpeak was the maximal cytosolic [Ca²⁺] within the beat.

#### Alternans magnitude and gain

ΔAPD and ΔCaTpeak were the beat-to-beat differences between the last two beats (99^th^ and 100^th^) at each CL. In single cells, the long beat was defined as the beat with the larger CaT, and ΔAPD and ΔCaTpeak were taken as magnitudes. Alternans onset was the longest CL at which ΔAPD > 1 ms (APD alternans) or ΔCaTpeak > 0.05µM (CaT alternans). Alternans gain was computed as ΔAPD/ΔCaTpeak (ms µM⁻¹) at each CL.

#### Electromechanical concordance

APD and CaT alternans were electromechanically concordant when the long APD coincided with the large CaT, and discordant otherwise.

#### Spatial concordance and discordance

In the strand, signed differences were used, so a sign change along the strand marks a phase reversal (nodal point). Analysis was restricted to cells 50-349 (0.625-4.375 cm) to exclude the stimulus site and sealed boundaries. At each CL, APD (or CaT) alternans was classified as: absent, if |ΔAPD| < 1 ms (|ΔCaT_peak| < 0.05 µM) at all analyzed cells; spatially concordant (SCA), if the central unstimulated region alternated in the same phase as the flanking stimulated regions. Spatially discordant (SDA), within the central unstimulated region reversed phase relative to the flanking regions. A tolerance of 0.1 ms (ΔAPD) or 0.005 µM (ΔCaTpeak) was used to reject numerical noise.

#### Severity of spatially discordant alternans

In-phase cells have the same sign of ΔAPD and out-of-phase cells have opposite signs. The phase was oriented such that ΔAPD was positive in the flanking stimulated regions. The severity of APD discordance was quantified as the most negative ΔAPD in the central unstimulated region at each CL. A larger negative value means that the central region alternates with a larger amplitude in antiphase with the flanking regions, and therefore a greater beat-to-beat spatial dispersion of repolarization.

#### Electrotonic entrainment

In the strand, electrotonic entrainment was defined as the synchronization of APD alternans in the central unstimulated region (ISO = 0 nM) with the flanking stimulated regions (ISO = 10 nM), driven by intercellular coupling currents that override the intrinsic alternans phase of the unstimulated region. The central region was considered entrained at a given CL when its ΔAPD had the same sign as the flanking regions, despite an opposite ΔCaT sign (i.e., APD concordance with CaT discordance).

### 2.5 Numerical implementation

The models were implemented in C++ (OpenMP-parallelized for the strand). The ionic model was integrated with a second-order Runge-Kutta scheme at a base time step of 0.01 ms, with automatic subdivision of the step when the local error estimate exceeded tolerance. Analysis was performed in MATLAB. All source code will be made available on GitHub at https://github.com/drgrandilab.

### 2.6 Optical mapping experiments

Optical mapping data from experiments described previously (10) were systematically re-analyzed to address aspects of spatial discordance of Ca^2+^/V_m_ alternans that were not previously assessed. All procedures were approved by the Animal Care and Use Committee of the University of California, Davis, and adhered to the NIH Guide for the Care and Use of Laboratory Animals.

Briefly, male New Zealand White rabbits were anesthetized (sodium pentobarbital, 50 mg kg⁻¹ IV). Hearts were excised and Langendorff-perfused at 37 °C. Hearts were stained with the voltage-sensitive dye, RH237 and loaded with the Ca²⁺ indicator, Rhod-2 AM. The excitation-contraction uncoupler, blebbistatin was used (10 µM). V_m_ and Ca²⁺ were recorded simultaneously with a dual-camera system (MiCAM Ultima, SciMedia/Brainvision; 100 × 100 pixels, 1 kHz). Hearts were paced from the LV basal epicardium with progressively shorter pacing cycle lengths (CL), from 300 ms to 120 ms. We analyzed CL = 140 ms, at which robust spatial alternans of both signals was present. Data presented here are from one heart at one CL and are presented descriptively; no statistical tests were performed.

### 2.7 Optical mapping analysis

Analysis was performed with custom MATLAB scripts. Each frame was spatially smoothed with a Gaussian filter (σ = 3 pixels). Each pixel trace was temporally smoothed with a 5-ms Gaussian window. Analysis was restricted to a manually drawn region of interest covering the mapped epicardium. Vm and CaT signals were normalized at every pixel. Activation time was calculated as 50% of the upstroke amplitude. APD70 was the interval from activation to 70% repolarization. APD alternans maps were generated by subtracting the even-beat APD70 from the odd-beat APD70, while CaT alternans maps were generated by subtracting the even-beat CaT peak from the odd-beat CaT peak. Nodal regions were pixels with |ΔAPD70| < 2.5 ms or |ΔCaT_peak| < 0.04 normalized units.

## 3. RESULTS

### 3.1 SR Ca²⁺ load alternation drives cellular alternans mediated by INaCa

Dynamic pacing of the baseline male endocardial cell model elicited sustained beat-to-beat alternation of both the CaT and APD (**Figure 1A**). Alternans remained beat-to-beat from onset down to CL 220 ms, after which higher-order alternans emerged before loss of capture at CL 200 ms. Alternans onset occurred at CL 260ms, with maximum ΔAPD (5.9 ms) and its corresponding ΔCaT (0.65 µM) at CL 245 ms. APD, peak CaT, ΔAPD, and ΔCaT vs. CL, computed from the last two beats at each CL, are shown in **Figure 1D**.

**Figure 1.**
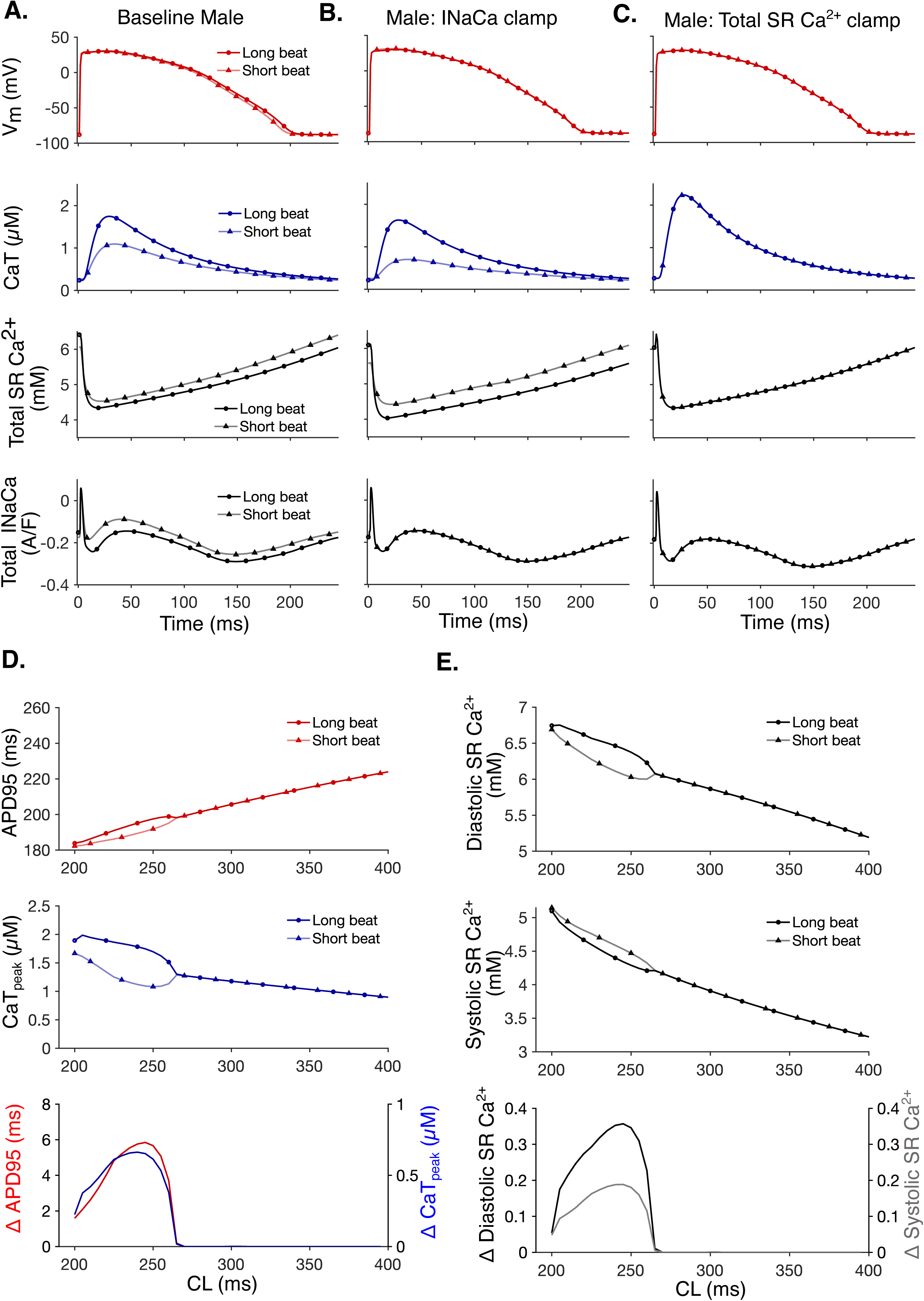
SR Ca²⁺ load alternation drives cellular repolarization alternans through INaCa. (**A)** Baseline male endocardial cell model under dynamic pacing. Time courses during dynamic pacing at CL = 245 ms: membrane potential (Vm, red), intracellular Ca²⁺ transient (CaT, blue), total SR Ca²⁺ content (Cansr+Cajsr, black), and total Na⁺/Ca²⁺ exchanger current (INaCa = I_NaCa,i + I_NaCa,ss, black), for the long beat (solid line, circles) and short beat (solid line, triangles), superimposed on a common time axis. Large CaT and SR Ca²⁺ content accompany the long APD and small CaT/SR content the short APD - Ca²⁺ and voltage alternans are in phase, indicating positive Ca²⁺→Vm coupling. **(B)** As in (A), but with total INaCa clamped to the long beat’s waveform. CaT alternans persist while APD alternans is abolished, showing that INaCa is required to translate Ca²⁺ alternans into APD alternans. **(C)** As in (A), but with total SR Ca²⁺ content (Cansr+Cajsr) clamped to the long beat’s waveform. Vm, Cai, SR Ca²⁺, and INaCa all converge to non-alternating waveforms, showing that clamping SR alone abolishes alternans entirely. **(D)** Restitution and alternans magnitude vs. cycle length, computed from the last two beats at each CL in the baseline male endocardial cell model. Top: APD95. Middle: peak CaT (Long beat: circles; short beat: triangles.) Bottom: beat-to-beat alternation magnitude, ΔAPD95 (red, left axis) and Δ CaTpeak (blue, right axis), vs. CL. Alternans onset at CL 260 ms; maximum ΔAPD95 (5.9 ms) and corresponding ΔCaT (0.65 µM) occur at CL 245 ms. **(E)** Diastolic (top) and systolic (middle) total SR Ca²⁺ content vs. CL (long beat: circles; short beat: triangles), and beat-to-beat SR alternation magnitude (bottom) in the baseline male endocardial cell model: Δdiastolic SR Ca²⁺ (black, left axis) and Δsystolic SR Ca²⁺ (gray, right axis) vs. CL. Both diastolic and systolic total SR Ca²⁺ alternate on every beat, indicating SR-load alternans in this model.

Consistent with prior work using the ORd family of models, these alternans were Ca²⁺-driven, whereby fluctuations in sarcoplasmic reticulum (SR) Ca²⁺ release and uptake produced the electrical APD alternation (42, 52). Large CaTs were associated with long APDs and small CaTs with short APDs, indicating positive Ca²⁺→Vm coupling. Coupling between CaT and APD alternans is mediated by the Na⁺/Ca²⁺ exchanger current (INaCa), consistent with human in vivo and in silico findings (52). Indeed, when clamping INaCa to the long-beat waveform (shown for CL 245 ms), CaT alternans persisted but APD alternans were abolished (**Figure 1B**). Both systolic and diastolic total SR Ca²⁺ (junctional plus network compartments) alternated between consecutive beats (**Figure 1E**), suggesting a load-dependent mechanism. In fact, clamping total SR Ca²⁺ to the long-beat waveform abolished alternans entirely (**Figure 1C**).

### 3.2 SERCA-dependent SR Ca²⁺ reuptake governs cellular alternans susceptibility

Because alternans are due to SR Ca^2+^ alternation in this model, we confirmed that altering SERCA shifted alternans susceptibility. Reducing SERCA activity by 40% exacerbated alternans, increasing maximum ΔAPD from 5.9 ms (baseline, **Figure 1D, 2A**) to 14 ms at CL 210 ms (**Figure 2C**).

**Figure 2.**
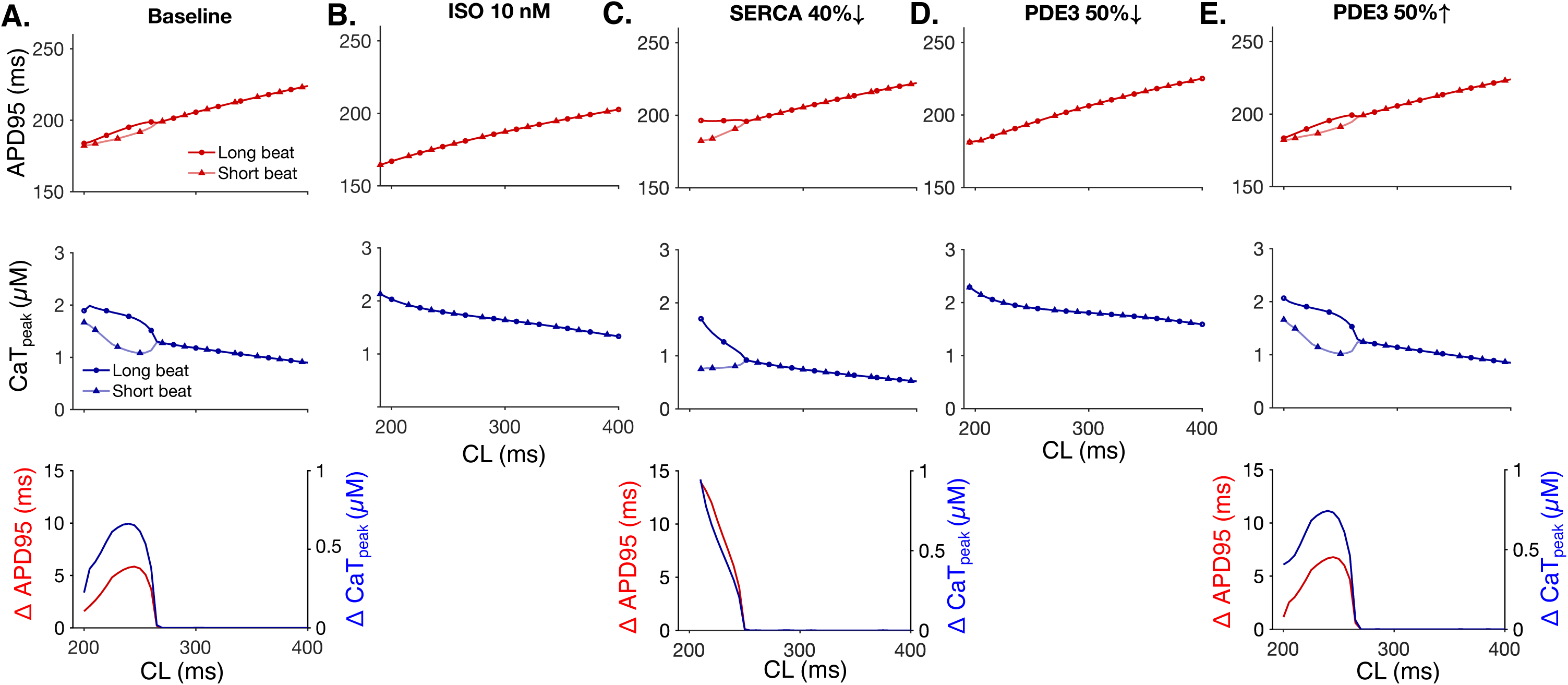
Enhanced SERCA-dependent SR Ca²⁺ reuptake suppresses repolarization alternans at the cellular level. In all panels: long beat - red circles (APD95) or blue circles (CaTpeak); short beat-triangles (APD95/CaTpeak panels, top two rows) computed from the last two beats at each CL; bottom row shows beat-to-beat alternation magnitude, ΔAPD95 (red, left axis) and Δ CaTpeak (blue, right axis), vs. BCL. (A) Baseline male endocardial cell model; (B) β-adrenergic stimulations with 10 nM ISO; (C) SERCA decreased by 40%; (D) PDE3 inhibition by 50%; (E) PDE3 enhancement by 50%;

Similarly, β-adrenergic stimulation with 10 nM ISO abolished both Ca²⁺ and APD alternans (**Figure 2B**), consistent with enhanced phospholamban phosphorylation and accelerated SR Ca²⁺ reuptake, as previously reported for β-adrenergic receptor stimulation in canine and human ventricular myocyte models (32) and in optically mapped rabbit hearts (10). PDE3 constrains the local cAMP signal that governs SERCA activity at the SR membrane (53). Manipulation of this mechanism of cAMP degradation produced corresponding effects, with PDE3 inhibition (×0.5) abolishing alternans entirely (**Figure 2D**) and PDE3 enhancement (×1.5) exacerbating it, increasing the maximum ΔAPD from 5.9 to 6.8 ms at CL 245 ms (**Figure 2E**).

### 3.3 Enhanced Ca²⁺-to-voltage coupling amplifies repolarization alternans in female myocytes

We next compared repolarization alternans susceptibility in the male and female ventricular myocyte models. The female model developed repolarization alternans at a longer CL than the male (280 versus 260 ms; **Figure 3A**), as previously shown (40), and female myocytes exhibited larger repolarization alternans, reaching a maximum ΔAPD of 8.2 ms at CL = 265 ms compared with 5.9 ms in the male at CL = 245 ms. Notably, the larger electrical alternans was not accompanied by larger Ca²⁺ alternans. At the CLs associated with maximum APD alternans, ΔCaTpeak was smaller in the female than in the male model (0.54 versus 0.65 µM). Thus, the sex difference in repolarization alternans could not be explained by the amplitude of the Ca²⁺ signal.

**Figure 3.**
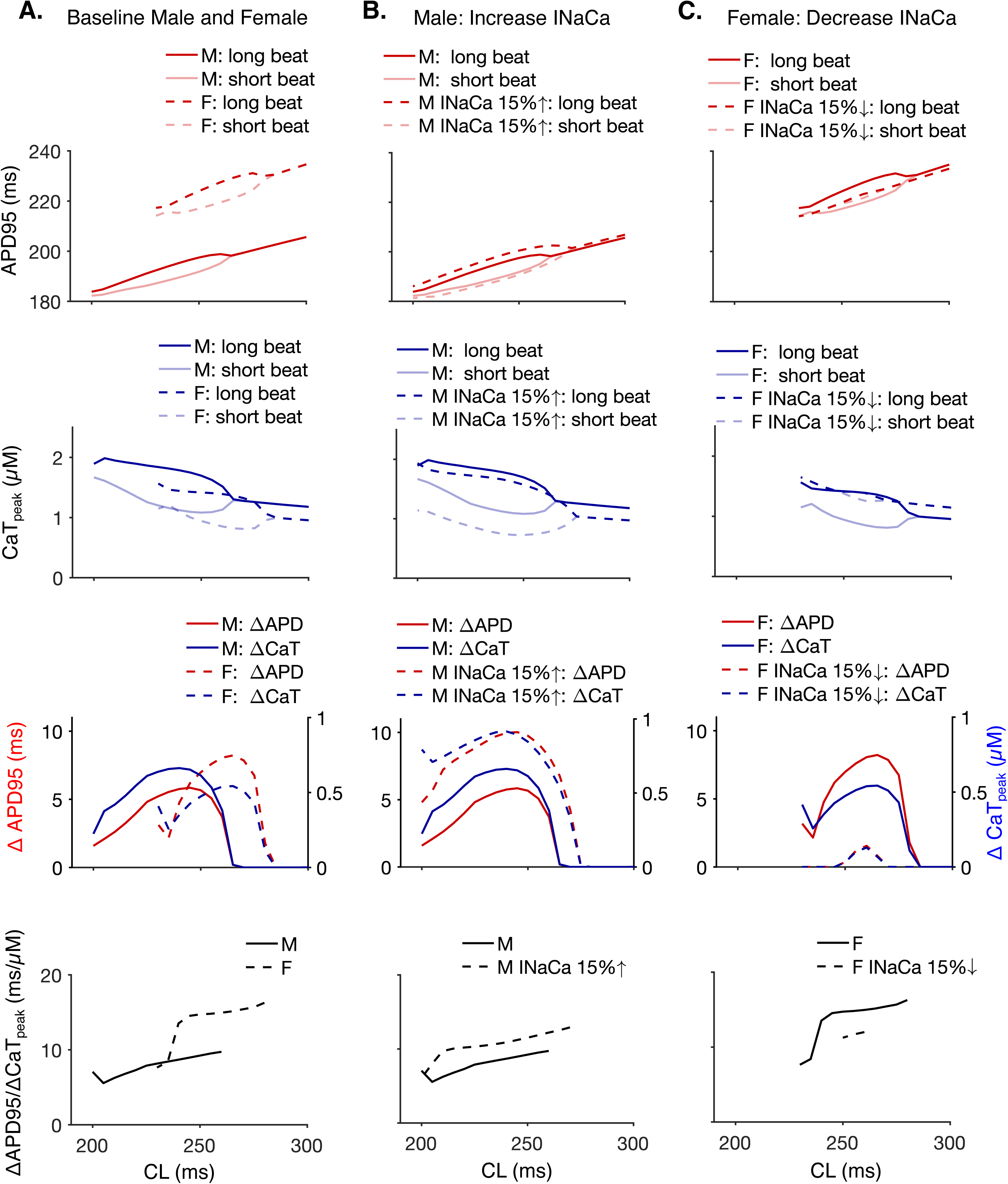
Higher INaCa-dependent Ca²⁺-to-voltage gain amplifies repolarization alternans in female myocytes. **A.** Male (solid line) vs. female (dashed) baseline model, BCL 200–300 ms. Top: APD95 restitution. 2nd row: CaTpeak restitution. 3rd row: ΔAPD (left axis) and ΔCaTpeak (right axis) vs. BCL. Bottom: alternans gain (ΔAPD/ΔCaTpeak, ms µM⁻¹) vs. BCL. **B.** Male baseline (solid) vs. male with INaCa scaled to the female level, INaCa×1.15 (dashed). Same row layout as A. **C.** Female baseline (solid) vs. female with INaCa scaled to the male level, INaCa×1 (dashed). Same row layout as A.

We therefore quantified the Ca^2+^-to-voltage coupling as the alternans gain, ΔAPD/ΔCaTpeak. The maximum gain was 1.7-fold higher in the female than in the male model (16.3 ms µM⁻¹ at CL 280ms versus 9.7 ms µM⁻¹ at CL 260; **Figure 3A**). Because the Na⁺/Ca²⁺ exchanger is the dominant route by which beat-to-beat differences in cytosolic Ca²⁺ are translated into differences in net repolarizing current (Results 3.1), and the female model has higher INaCa conductance as informed by previous human and animal studies reporting higher INaCa expression/activity in female vs. male myocardium (47, 54, 55), we tested its contribution to the gain sex difference.

Increasing INaCa in the male model to the female level increased maximum ΔAPD, ΔCaTpeak, and the ΔAPD/ΔCaT gain (5.9 to 10 ms, 0.65 to 0.9 µM, 9.7 to 12.9 ms µM⁻¹ ). Conversely, reducing INaCa in the female model to the male level decreased maximum ΔAPD from 8.2 to 1.6 ms, ΔCaT from 0.54 to 0.13 µM, and gain from 16.3 to 12.0 ms µM⁻¹ (**Figure 3B,C**). INaCa is therefore an important target for moving alternans gain in either direction.

Because altering INaCa also affects cellular Ca²⁺ balance, we next matched the Ca²⁺ signal between sexes. The female cytosolic Ca²⁺ waveform at CL = 245 ms was clamped to the male recorded waveform at the same CL, so that the two models experienced Ca²⁺ transients of identical amplitude and identical alternans magnitude (ΔCaTpeak = 0.65 µM in both). A male self-clamp control, driven by its own waveform under the same protocol, reproduced the male’s ΔAPD exactly (5.9 ms). Under the matched Ca²⁺ clamp, the female model still developed the larger repolarization alternans (8.9 versus 5.9 ms; not shown). Thus, female myocytes convert a given Ca²⁺ alternans into a larger repolarization alternans compared to male myocytes, demonstrating a sex difference in the Ca^2+^-to-voltage coupling downstream of the CaT.

### 3.4 Spatial heterogeneity in β-AR signaling affects spatial concordance of Ca^2+^ and repolarization alternans

We next examined how spatial heterogeneity in β-adrenergic signaling affects alternans in a one-dimensional strand. A 5-cm strand was exposed to 10 nM isoproterenol (ISO) except for a central 0.3-cm region with no ISO, representing a localized region of reduced sympathetic stimulation (**Figure 4A**, 1^st^ row). The strand was paced dynamically from one end.

**Figure 4.**
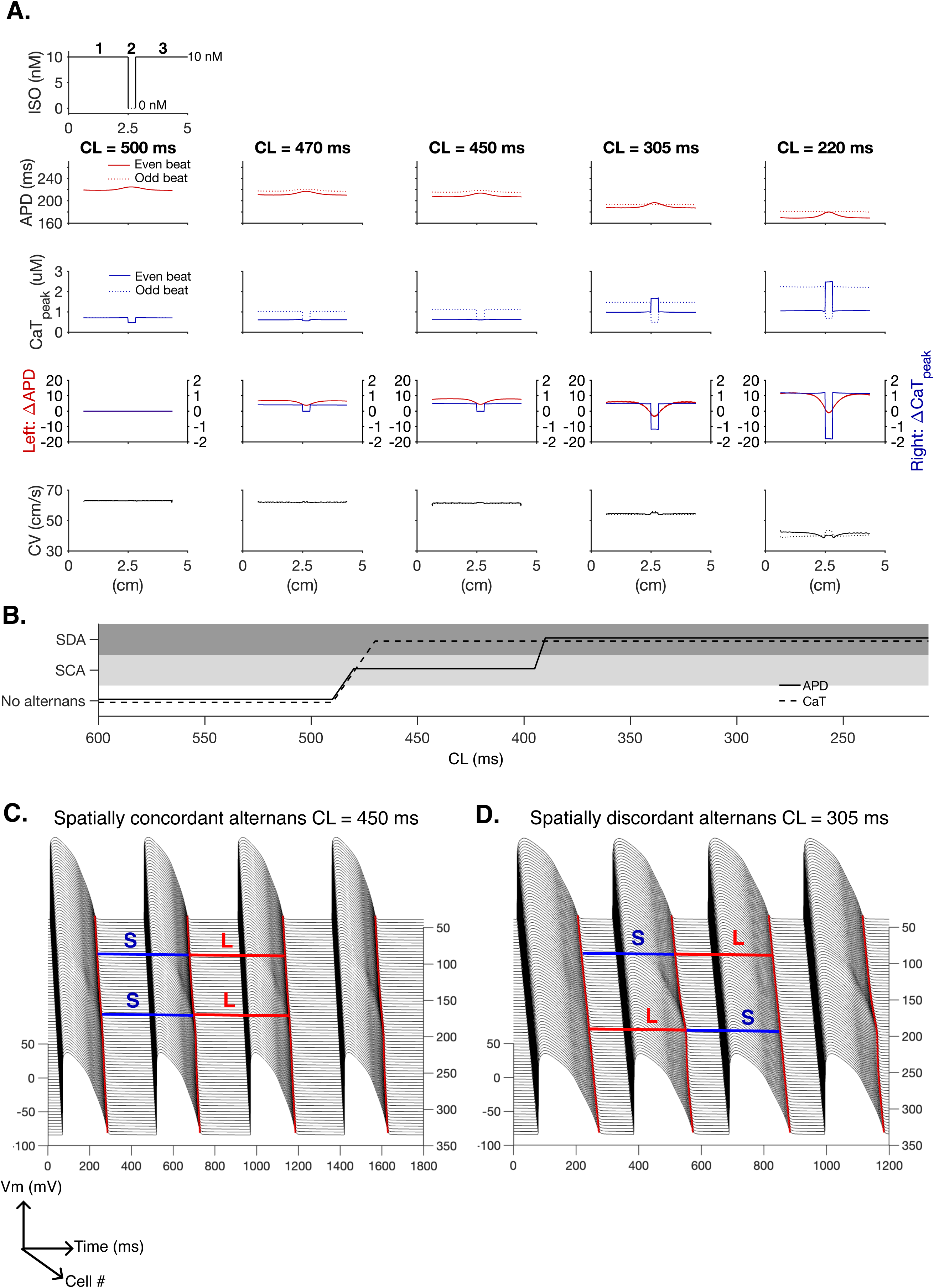
Spatial heterogeneity in β-AR signaling affects spatial concordance of Ca^2+^ and repolarization alternans. **A.** Row1: Spatial ISO exposure profile along the 1D strand: background ISO = 10 nM in regions 1 and 3, flanking a central 0.3 cm patch of ISO = 0 nM. APD95 (Row 2), CaTpeak (Row 3), ΔAPD (left axis) and ΔCaTpeak (right axis) (Row 4), and conduction velocity (CV) (Row 5), vs. fiber position (cm) computed from the last two beats at five representative cycle lengths (CL = 500, 470, 450, 305, and 220 ms); **B.** Alternans classification (no alternans / spatially concordant alternans, SCA / spatially discordant alternans, SDA) vs. pacing cycle length (CL), determined from behavior within the ISO = 0 patch (region 2), shown independently for APD (solid) and CaTpeak (dashed). Representative spatiotemporal Vm maps illustrating **C.** spatially concordant alternans (SCA, left, CL = 450 ms) vs **D.** spatially discordant alternans (SDA, right, CL = 305 ms). L/S labels mark the long (L) and short (S) beat in each region; under SCA the same beat identity (L or S) is longest or shortest everywhere along the fiber, whereas under SDA region 2’s long beat coincides in time with regions 1 and 3’s short beat, and vice versa.

The onset and magnitude of CaT alternans depend on the balance between SR Ca²⁺ release and reuptake and is set by diastolic SR Ca²⁺ load, the steepness of the release-load relationship, the SERCA uptake rate, RyR2 recovery, and the ICaL trigger (32, 52, 56, 57). Each of these is altered by β-AR signaling. In baseline male and female strands, pacing from one end did not produce alternans in the middle of the strand, unlike single-cell pacing at the same cycle length. This reflects a difference between the Ca²⁺-release trigger delivered by a propagating action potential versus a clamped/isolated single-cell stimulus, as previously described by (32), who showed that cell-to-cell coupling blunts the AP upstroke and thereby reduces the trigger for SR Ca²⁺ release in 1D strand simulations relative to single cells. When all cells in the strand were stimulated simultaneously instead of paced from one end, alternans appeared and reproduced single-cell behavior (not shown).

To obtain robust alternans under strand pacing necessary for studying its spatial patterning SERCA uptake was reduced by 40% throughout the strand, mimicking a heart-failure like phenotype and consistent with reduced SR Ca²⁺-handling reserve promoting alternans (16–18). All simulations described below use this reduced-SERCA background. With 40% SERCA reduction, 10 nM ISO is insufficient to abolish repolarization alternans throughout the cable.

As shown in the male endocardial strand, at long CLs, the unstimulated central region exhibited longer APD and smaller CaT amplitude than the surrounding ISO-stimulated regions, as expected from the absence of β-AR/PKA-dependent stimulation of L-type Ca²⁺ current, SR Ca²⁺ release, IKs, and SERCA (**Figure 4A,B, CL 500ms**). At increasing pacing rates, CaT alternans became spatially discordant first, with phase reversal in the central region appearing at CL = 470 ms. APD alternans remained spatially concordant at this CL because electrotonic coupling smooths voltage over a much longer length scale than the underlying Ca²⁺ instability: CaT alternans can reverse phase over roughly one cell length (27), while APD alternans reverses phase only over the length scale set by electrotonic coupling, as established by (25, 26). As pacing CL was reduced further and CaT discordance grew, APD alternans also became spatially discordant at CL = 390 ms. Nodal (crossover) points became visible in APD in the patch, with the maximum APD discordance (ΔAPD = −3.6 ms) occurring at CL = 305 ms (**Figure 4B,C, CL 305ms**). The ISO= 0 and ISO-stimulated regions of the strand operate at different SR Ca²⁺ loads, so at a given CL the two regions have different Ca²⁺ dynamics.

Whether the unstimulated region develops spatially discordant APD alternans depends on the interplay of several factors: the amplitude of CaT discordance itself, the strength of electrotonic coupling, the Ca²⁺-to-APD coupling gain, and the magnitude of APD alternans already present in the ISO-stimulated regions. For example, at CL = 220 ms the magnitude of APD discordance (negative ΔAPD =-1.2ms) in the unstimulated region decreased relative to intermediate CLs despite CaT alternans remaining strongly discordant: ΔAPD in the surrounding ISO-stimulated regions is large (11-12 ms) and pulling the central region toward the same alternating phase as the now strongly-alternating flanking regions.

CV showed little change across cells at the longer CLs where CaT/APD discordance first appeared (**Figure 4A, B**), suggesting that CV restitution is not the dominant driver of discordance onset. Instead, discordance originated from the underlying ISO-driven functional heterogeneity between the central unstimulated region and the surrounding ISO-stimulated areas. This is consistent with computational work showing that when CV restitution is not strongly engaged, spatially discordant alternans can still form at and remain pinned to pre-existing tissue heterogeneities, rather than requiring restitution-driven instability (58). At shorter CL (e.g., 220 ms), however, beat-to-beat changes in conduction contributed to the spatial pattern: at a given beat, a short diastolic interval (DI), reduced Na⁺ channel recovery and slowed conduction on the subsequent beat, lengthening the downstream DI and APD, while a long DI accelerated conduction and shortened the downstream DI and APD - the classical DI-restitution feedback loop implicated in CV-driven discordant alternans (8, 59). In a strand with uniformly reduced SERCA but spatially uniform ISO (either 0 or 10 nM throughout), spatial alternans remained concordant in both APD and CaT (*not shown*), confirming that spatially discordant alternans in this model specifically required the imposed ISO heterogeneity rather than arising spontaneously from CV restitution alone.

### 3.5 Higher Ca^2+^ to voltage gain limits APD discordance in the female strand

We next repeated the heterogeneous ISO protocol in the female strand using the same geometry and 40% reduction in SERCA activity (**Figure 5A** row1). Given the greater Ca²⁺-to-Vm coupling gain observed in female cells, we initially expected greater susceptibility to spatially discordant APD alternans. Instead, the female strand exhibited substantially less APD discordance than the male strand. CaT alternans in the central region became spatially discordant at CL = 460 ms, yet APD alternans remained concordant over most of the pacing range. APD discordance appeared only between CL = 325 and 250 ms and remained small, with a maximum negative ΔAPD of -0.8 ms (compared to -3.6 ms under the same spatial ISO profile).

**Figure 5.**
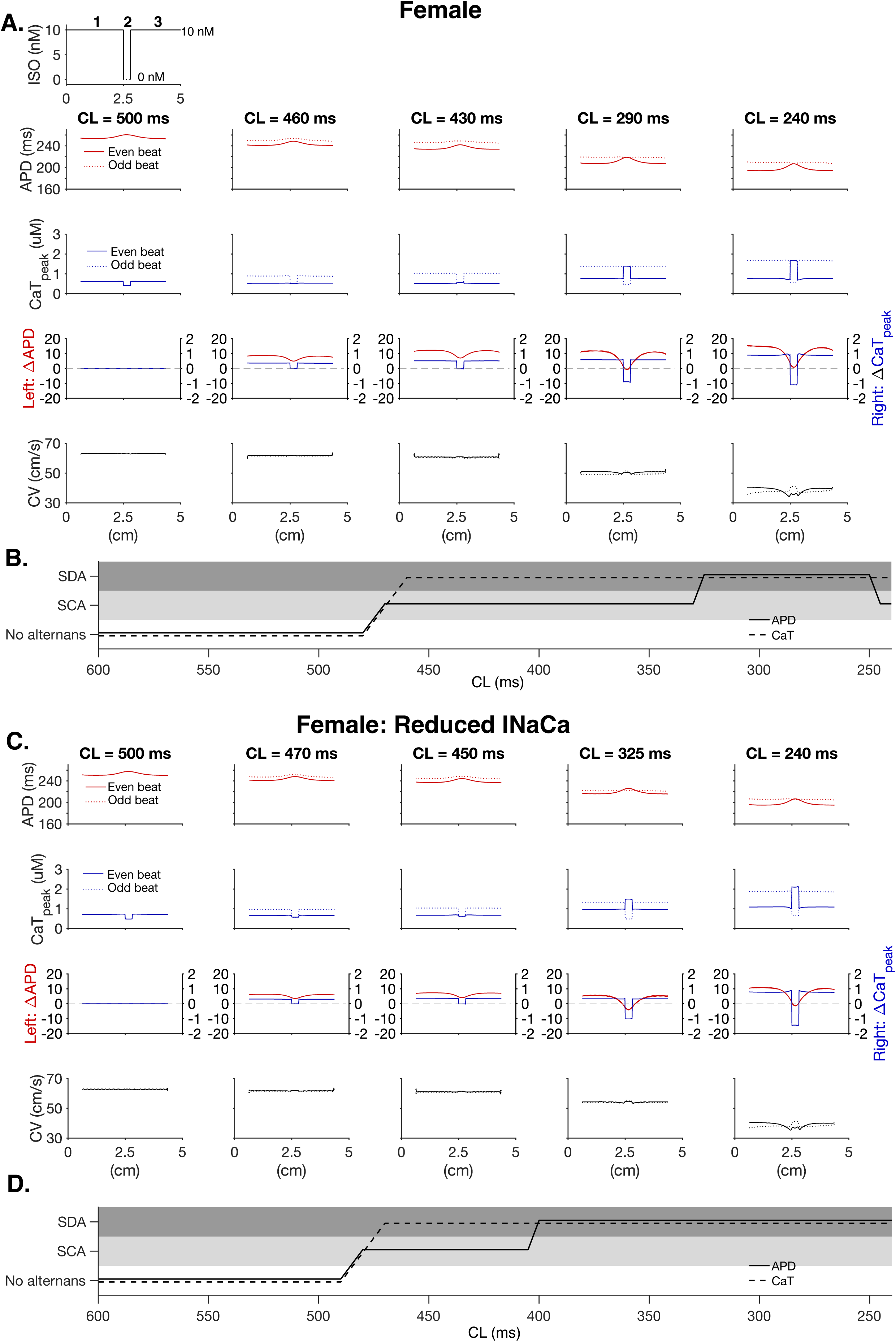
Higher Ca^2+^-to-voltage coupling limits APD discordance in the female ventricular strand. **A.** Row1: Spatial ISO exposure profile along the 1D strand: background ISO = 10 nM in regions 1 and 3, flanking a central 0.3 cm patch of ISO = 0 nM. APD95 (Row 2), CaTpeak (Row 3), ΔAPD (left axis) and ΔCaTpeak (right axis) (Row 4), and conduction velocity (CV) (Row 5), vs. fiber position (cm) computed from the last two beats at five representative cycle lengths (CL = 500, 460, 430, 290, and 240 ms) for the female strand with baseline INaCa; **B.** Alternans classification (no alternans / spatially concordant alternans, SCA / spatially discordant alternans, SDA) vs. cycle length (CL), determined from behavior within the ISO = 0 patch (region 2), shown independently for APD (solid) and CaTpeak (dashed). **C**. and **D.** Same layout as **A.** and **B.**, for the female strand with INaCa reduced to the baseline level in the male model.

This finding can once again be explained by the greater Ca²⁺-to-Vm gain in the female model. The large ΔAPD alternans produced by the high gain in the ISO-stimulated regions (flanking the ISO = 0 patch) electrotonically entrains the central unstimulated region into the same alternating phase, preserving APD concordance even when local Ca²⁺ transient is discordant (**Figure 5A, B**). This is consistent with the general principle that the phase of a spatially patterned alternans instability is set by the interplay between the local instability and the amplitude/coupling of the surrounding tissue (60). CaT alternans in the unstimulated region became spatially discordant at CL = 460 ms. When the pacing CL was reduced between 325 and 250 ms, the region eventually crossed into small APD discordance, but the effect remained small: the maximum negative ΔAPD did not exceed −0.8 ms across this range. In the male strand, by contrast, the equivalent discordance reached as negative as −3.6 ms (at PCL = 305 ms).

To test this mechanism directly, we reduced INaCa in the female strand to the male level. This manipulation markedly increased susceptibility to APD discordance: phase reversal developed at longer CLs and reached −4.2 ms at CL = 325 ms (**Figure 5C,D**), similar to the behavior of the male strand.

The results show that the stronger INaCa-dependent Ca²⁺-APD coupling in the female model enforces global APD phase-locking that overrides locally discordant Ca²⁺ signaling and limits the translation of local Ca^2+^ discordance into APD discordance.

### 3.6 Reduced electrotonic coupling enhances spatially discordant alternans susceptibility in both sexes

If APD concordance is maintained by electrotonic entrainment, weakening intercellular coupling should allow the local Ca²⁺ instability to exert greater control over APD phase. We therefore repeated the heterogeneous ISO protocol after reducing electrotonic coupling uniformly across the strand in both sexes.

In the male model, reducing coupling shifted the onset of APD discordance to a longer cycle length, from 390 ms with normal coupling (**Figure 4B**) to 430 ms with reduced coupling. The magnitude of discordance was also substantially larger under weak coupling, reaching −9.3 ms at CL = 340 ms (**Figure 6A**), compared with −3.6 ms at normal coupling. CaT alternans discordance appeared at CL = 490 ms under weak coupling, again preceding APD discordance. The female model, which was largely protected from SDA at normal coupling, lost that protection under weak coupling: APD discordance now appeared at CL = 405 ms, and its magnitude reached −9.7 ms at CL = 285 ms (**Figure 6C,D**), while CaT alternans discordance began at CL = 480 ms under weak coupling. Thus, under reduced coupling, male and female strands developed similarly large APD phase reversals.

**Figure 6.**
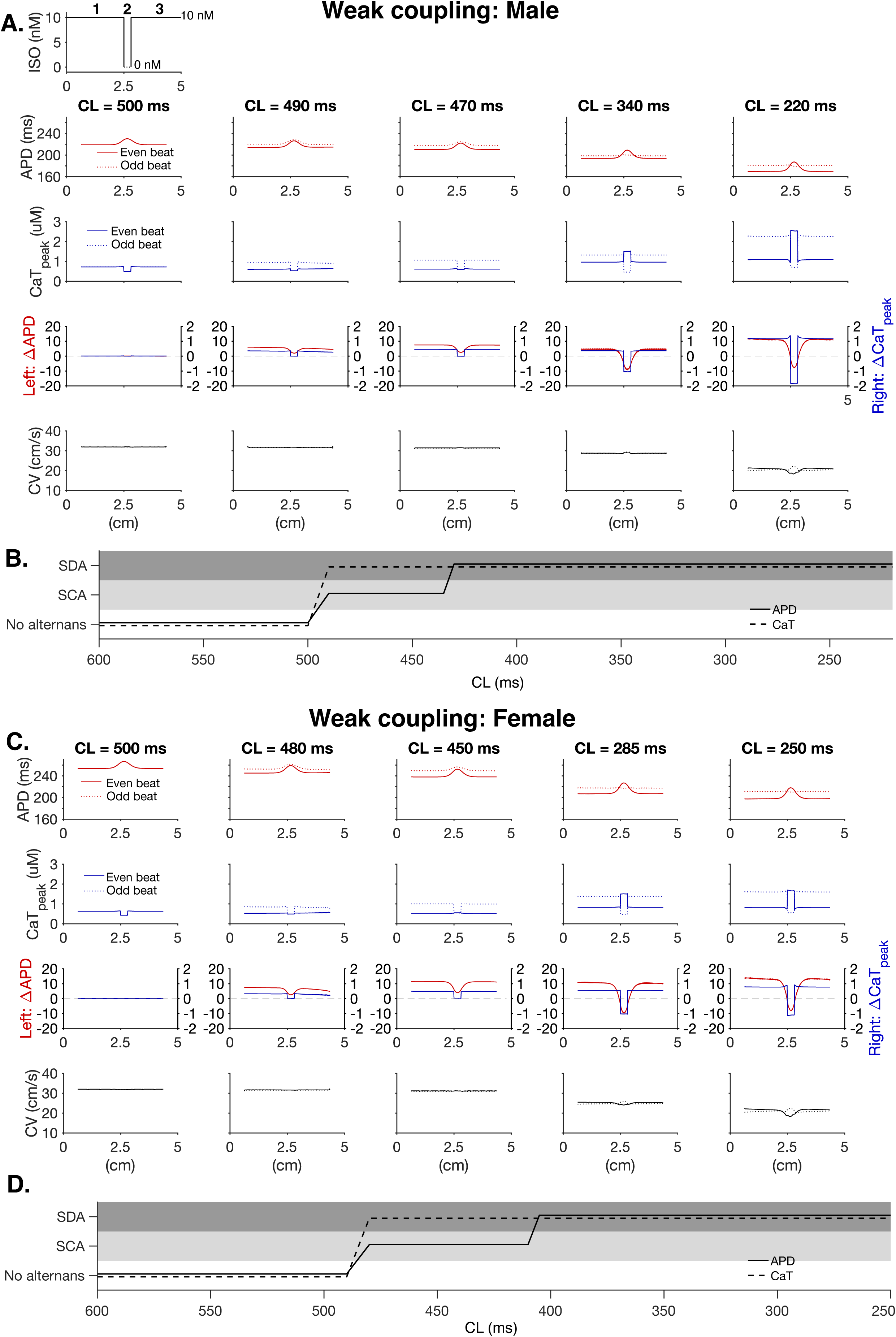
Reduced electrotonic coupling increases susceptibility to spatially discordant alternans in both sexes. **A.** Row1: Spatial ISO exposure profile along the 1D strand: background ISO = 10 nM in regions 1 and 3, flanking a central 0.3 cm patch of ISO = 0 nM. APD95 (Row 2), CaTpeak (Row 3), ΔAPD (left axis) and ΔCaTpeak (right axis) (Row 4), and conduction velocity (CV) (Row 5), vs. fiber position (cm) for the male strand under reduced electrotonic coupling, computed from the last two beats at five representative cycle lengths (CL = 500, 490, 470, 340, and 220 ms) **B.** Alternans classification (no alternans / SCA / SDA) vs. PCL for the male strand under reduced coupling, APD (solid) and CaTpeak (dashed); **C.** Same layout as A., for the female strand under reduced electrotonic coupling, at five representative CLs (500, 480, 450, 285, and 250 ms). **D.** Same layout as B., for the female strand under reduced electrotonic coupling.

### 3.7 Enlarging the heterogeneous ISO void region promotes SDA in both sexes

Electrotonic entrainment should also depend on the spatial extent of the heterogeneous region, independent of any change in coupling strength itself, because what ultimately matters is the size of the heterogeneous region relative to the electrotonic length constant over which APD alternans phase is smoothed (26, 29). We tested this directly by doubling the width of the ISO = 0 patch from 0.3 to 0.6 cm, keeping normal electrotonic coupling and all other parameters identical to Figure 4.

In the male strand, enlarging the patch shifted APD discordance onset from CL = 390 ms to CL = 415 ms, and increased the peak discordance magnitude from -3.6 ms to -9.5 ms at CL = 305 ms (**Figure 7A, B**). CaT alternans discordance appeared at CL = 470 ms under the enlarged patch. In the female strand, the shift was particularly pronounced: while APD discordance was marginal with the 0.3 cm unstimulated region (max -0.8 ms, over CL 325–250 ms; Results 3.5), increasing the width to 0.6 cm produced APD discordance at CL = 385 ms, with peak magnitude reaching - 9.5 ms at CL = 260 ms (**Figure 7C, D**). CaT alternans discordance appeared at CL = 460 ms.

**Figure 7.**
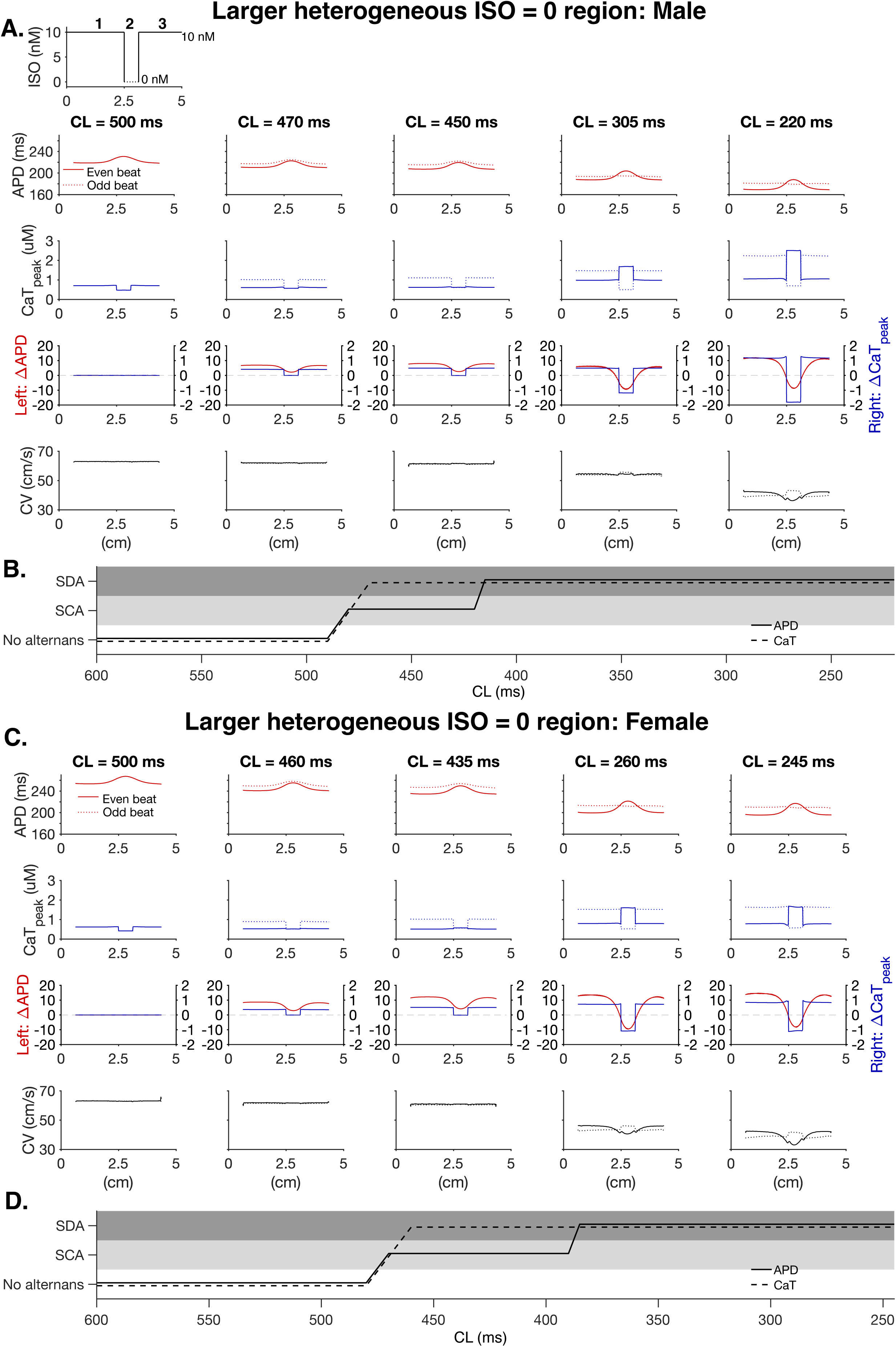
Spatial extent of β-adrenergic heterogeneity determines susceptibility to APD discordance. **A.** Row1: Spatial ISO exposure profile along the 1D strand: background ISO = 10 nM in regions 1 and 3, flanking a central 0.6 cm patch of ISO = 0 nM. APD95 (Row 2), CaTpeak (Row 3), ΔAPD (left axis) and ΔCaTpeak (right axis) (Row 4), and conduction velocity (CV) (Row 5), vs. fiber position (cm) for the male strand with an enlarged (0.6 cm) ISO = 0 region, computed from the last two beats at five representative cycle lengths (CL = 500, 470, 450, 305, and 220 ms) **B.** Alternans classification (no alternans / SCA / SDA) vs. PCL for the male strand with the enlarged ISO = 0 region, APD (solid) and CaTpeak (dashed); **C.** Same layout as A., for the female strand with the enlarged ISO = 0 region, at five representative CLs (500, 460, 435, 260, and 245 ms). **D.** Same layout as B., for the female strand with the enlarged ISO = 0 region.

Thus, enlarging the heterogeneous region and weakening electrotonic coupling produced qualitatively similar effects: both reduced the ability of the flanking regions to impose their APD phase on the central region and narrowed the difference between male and female strands. The ratio of heterogeneity size to the electrotonic length constant (itself set by coupling strength) determines whether the flanking tissue can impose its phase on the central region (26, 29). Once this ratio exceeds a critical value, whether by shrinking the length constant or enlarging the heterogeneous region, the female’s higher Ca²⁺-to-APD coupling gain is no longer sufficient to enforce phase-locking, and both sexes converge on comparable SDA severity.

### 3.8. Optical mapping of the rabbit ventricle shows APD phase locking that preserves APD concordance across a narrow region of CaT discordance

To test whether the APD phase locking identified in the strand simulations (Results 3.4 -3.7) operates in intact myocardium, we analyzed simultaneous epicardial Vm and Ca²⁺ optical maps from a Langendorff-perfused healthy male rabbit heart with a dynamic protocol of decreasing CL (see Methods). We focused on CL = 140 ms, at which robust alternans were present in both signals and a region of spatially discordant CaT alternans coexisted with spatially concordant APD alternans (**Figure 8**).

**Figure 8.**
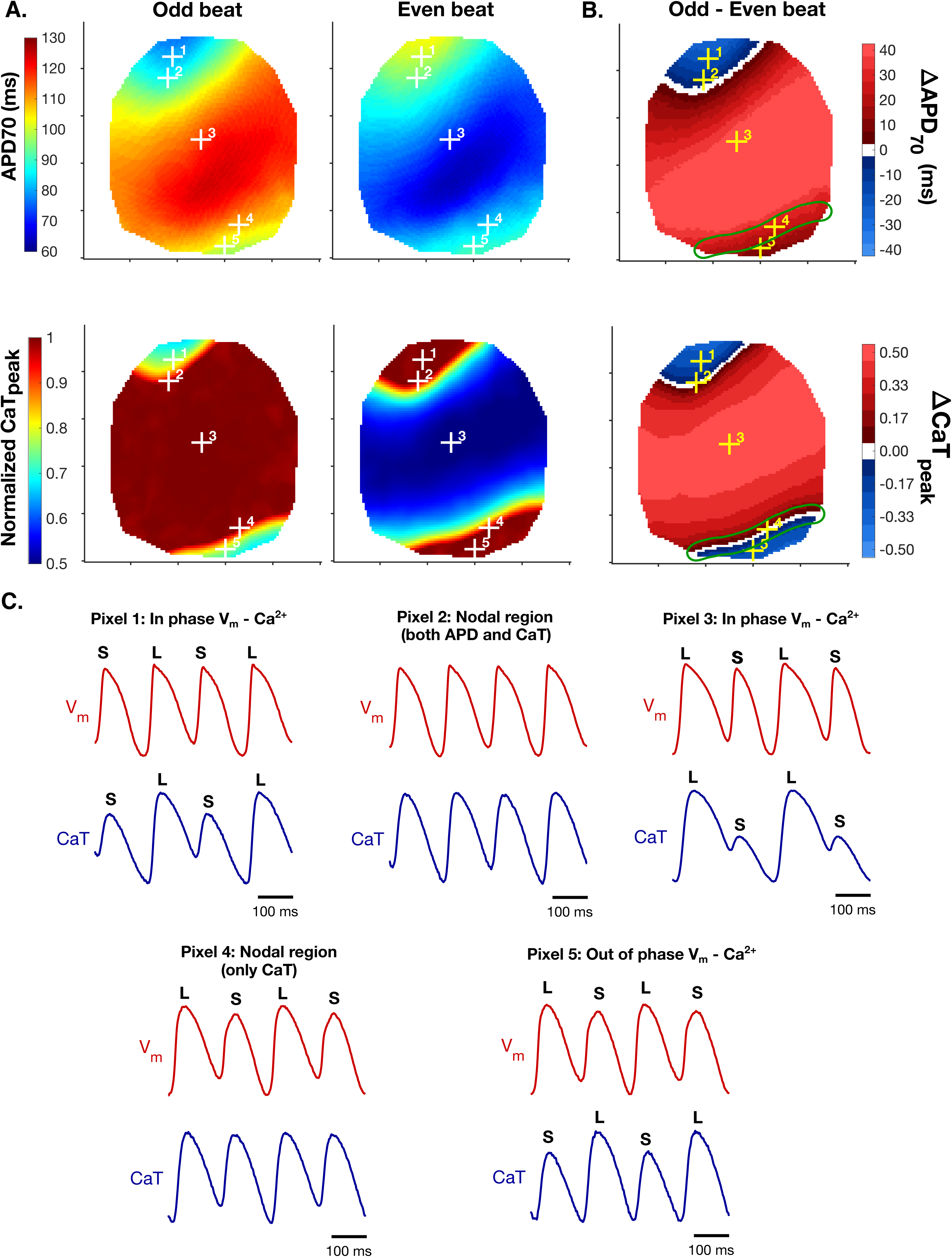
Optical mapping of the rabbit ventricle shows APD phase locking that preserves APD concordance across a narrow region of CaT discordance. Simultaneous epicardial Vm and Ca²⁺ optical maps from a Langendorff-perfused healthy male rabbit heart paced at CL = 140 ms. Vm and CaT signals were normalized at every pixel. **(A)** APD70 maps (top row) and CaTpeak maps (bottom row) for an odd and the following even beat. **(B)** Beat-to-beat difference maps (odd - even beat) of ΔAPD70 (top) and ΔCaTpeak (bottom). Red and blue mark the two opposite alternans phases, and white marks near-zero alternans, i.e. nodal lines (|ΔAPD70| < 2.5 ms; |ΔCaT_peak| < 0.04). The green outline encloses the lower CaT nodal region and is superimposed at the same position on the ΔAPD70 map; in this region CaT reverses phase while APD keeps the phase of the central region. **(C)** Vm (red) and CaT (blue) traces over four consecutive beats at pixels 1-5 (marked + in A and B). L and S label the long and short APD, and the large and small CaT, of each beat. Pixel 1: Vm and CaT in phase, but alternating opposite to the central region. Pixel 2: adjacent to the common APD and CaT nodal line. Pixel 3: central region with large in-phase alternans. Pixel 4: on the CaT nodal line only (APD alternans remains large). Pixel 5: Vm and CaT out of phase (CaT discordant, APD concordant with the central region). Scale bars: 100 ms.

Most of the mapped surface alternated in a single phase, with the largest alternans in the central region (pixel 3: ΔAPD70 = 44 ms, ΔCaT_peak = 0.5; **Figure 8B, C**). Here APD and CaT alternated in phase (long APD with large CaT), consistent with the positive Ca²⁺ to Vm coupling. The central region was bordered by two regions in which CaT alternated in the opposite phase, and these differed in whether APD followed.

In the upper region, APD and CaT reversed phase together. Pixel 1 alternated in phase electromechanically (short APD with small CaT); but out of phase with the central region, i.e., both APD and CaT alternans were spatially discordant. The two signals shared a common nodal region: at pixel 2, which adjacent to the common line, ΔCaTpeak was ∼0 and ΔAPD70 was -7 ms, with APD alternans in its vicinity within ±5 ms.

In the lower region, by contrast, CaT reversed phase but APD did not. Pixel 4 lies on the CaT nodal line, yet APD alternans there remained large and in phase with the central region (ΔAPD70 = 20 - 30 ms along the CaT nodal line, green outline in **Figure 8B**). Within the narrow region of CaT discordance beyond it (pixel 5), CaT alternated out of phase with the central region while APD retained the central phase, so that Vm and CaT alternated out of phase locally (long APD with small CaT).

These observations reproduce the behavior of the simulated strands. CaT alternans reversed phase over a short distance, whereas APD phase was smoothed over the longer length scale set by electrotonic coupling (25, 26). Whether APD followed the CaT reversal depended on the two factors identified *in silico*. In the lower region, the CaT-discordant band was narrow and bordered by tissue with large APD alternans ΔAPD70 of 20-45 ms, which electrotonically entrained repolarization and prevented APD phase reversal, analogous to phase locking by high-ΔAPD flanking regions in the strand (Results 3.4, 3.5). In the upper region, APD alternans near the nodal line was small allowing APD to reverse with CaT.

## 4. DISCUSSION

### 4.1 Summary of Findings

Using sex-specific human ventricular myocyte models incorporating β-adrenergic signaling, coupled in one-dimensional strands containing a localized region lacking β-AR stimulation, we identify a scale-dependent sex difference in repolarization alternans. At the cellular level, female myocytes develop larger APD alternans despite smaller CaT alternans than male myocytes, reflecting higher Ca²⁺-to-voltage gain. This difference is INaCa-dependent and persists when the Ca²⁺ transient is matched between sexes, demonstrating that it arises downstream of Ca²⁺ cycling itself. In a coupled strand, however, the larger APD alternans in β-AR-stimulated regions electrotonically entrains the unstimulated region, maintaining APD phase-locking despite locally discordant CaT alternans. In contrast, the male strand develops substantial APD discordance under the same βARS heterogeneity. Optical maps of the rabbit ventricle demonstrate qualitatively similar, physiologically relevant patterns, with a narrow band of discordant CaT alternans bordered by large-ΔAPD tissue in which APD retains the surrounding phase.

Two aspects are, to our knowledge, particularly notable and novel. First, the greater susceptibility of female myocytes to repolarization alternans reflects enhanced Ca²⁺-to-voltage coupling, primarily associated with greater INaCa. Second, and more consequentially, the functional consequence of this higher gain changes with spatial scale: it amplifies APD alternans in isolated cells but can stabilize APD phase in coupled cells by strengthening electrotonic entrainment.

### 4.2 Sex differences and clinical implications

Clinically, women have lower rates of sustained VT/VF and SCD than men (38, 39) despite longer QT, reduced repolarization reserve and greater susceptibility to drug-induced torsades de pointes (61–63). Females also show greater repolarization sensitivity to β-AR stimulation, with larger transient QTc prolongation and more frequent bifid T-wave morphology (64), and steeper HR-QT relations under adrenergic stimulation (65). Longer intrinsic QTc persists under autonomic blockade, indicating intrinsic electrophysiological sex differences independent of autonomic tone (41, 47, 66). Our results may offer a mechanistic explanation of this pattern. A higher ΔAPD for a given ΔCaT would be expected to amplify repolarization alternans when alternans remain spatially concordant, consistent with greater repolarization lability in females. Once Ca alternans become spatially discordant, however, the same high Ca-to-voltage gain strengthens electrotonic APD phase-locking, limiting the repolarization gradients that favor reentry. This may help reconcile the greater female susceptibility to some forms of repolarization instability with the lower incidence of reentrant VT/VF and SCD, although this possibility requires direct experimental testing.

The findings also have implications for sympathetic modulation. Several studies report β-blockade reduces TWA incidence and alternans magnitude in patients (67, 68), consistent with the well-established anti-arrhythmic effects of reducing adrenergic drive, heart rate, and triggered activity (69). Our results suggest that the spatial distribution of β-adrenergic signaling (30) may contribute to the net outcome. While β-AR stimulation suppressed alternans at the single-cell level, spatially nonuniform stimulation created regional differences in alternans susceptibility that could promote Ca²⁺ and APD phase discordance. Thus, therapies that reduce regional gradients in sympathetic influence may provide benefits beyond simply lowering global adrenergic tone, particularly in hearts with heterogeneous denervation or hyperinnervation. More broadly, the relationship between TWA amplitude and the underlying spatially discordant substrate may depend on how adrenergic signaling is distributed across the myocardium.

Our findings also suggest that this relationship may be sex-dependent. Higher INaCa expression and activity in female myocardium has been reported across human and animal preparations (47, 54, 55, 70), and in our model this contributed to higher Ca²⁺-to-voltage coupling gain. Interventions acting on Ca²⁺ cycling stability (e.g., SERCA activators, RyR2 stabilizers) may be less protective in female vs. male myocytes for a given improvement in Ca²⁺ handling, whereas targeting INaCa may be more effective. Consistent with a role for INaCa, the selective INaCa blocker ORM-10962 has been shown to attenuate both APD and CaT alternans (71).

The reduced susceptibility of the female strand to APD discordance was lost both when electrotonic coupling was weakened and when the unstimulated region was enlarged, with male and female strands converging toward similarly large SDA. Both manipulations reduce the ability of the surrounding myocardium to impose its APD phase on the heterogeneous region. Conceptually, they increase the size of the heterogeneous region relative to the electrotonic length scale over which voltage is smoothed (25, 26, 29). As this ratio increases, local Ca²⁺ dynamics exert greater control over APD phase and electrotonic phase-locking becomes less effective. This interpretation is consistent with previous work identifying intercellular coupling as a major determinant of discordant-alternans onset and domain size (23), and with evidence that preserving Cx43-mediated coupling suppresses discordant alternans and VT/VF rearrest in vivo (24). Therefore, the reduced SDA susceptibility in female strands requires strong electrotonic coupling and a heterogeneous region small enough to remain electrotonically entrained. Disease-associated gap-junction remodeling, fibrosis, or larger regions of sympathetic heterogeneity would therefore be expected to diminish this sex difference.

### 4.3 Limitations and future directions

Alternans in this model rely on SR Ca²⁺ load alternation and are not sustained by release refractoriness alone (9, 10, 72). This differs from isolated ventricular myocytes (73) and the intact rabbit heart (10), where SR Ca²⁺ release alternans arises without any change in diastolic SR Ca²⁺ at longer cycle lengths and load contributes at shorter CL. The present findings should be interpreted within the Ca²⁺-cycling properties of this model, whereas reproducing refractoriness-driven alternans in the ORd/Heijman-Rudy family requires explicit modification of SR release dynamics (57) that could be done in future studies.

The multicellular simulations were limited to one-dimensional strands, which cannot support reentry. We therefore infer arrhythmogenic potential from the emergence of a SDA substrate. Future work will extend this framework to 2D simulations to quantify how β-AR heterogeneity and sex differences interact to shape alternans dynamics and arrhythmia vulnerability.

The onset and spatial pattern of alternans depend not only on the instantaneous CL but also on pacing history. Slowly accumulating state variables, such as intracellular Na⁺ and SR Ca²⁺, introduce short-term memory, making restitution and alternans thresholds protocol-dependent (74). In this study, we used a gradual dynamic pacing protocol, decreasing CL in small increments and allowing 100 beats at each CL for beat-to-beat transients to settle before the next decrement. Within the alternans regime, CL was reduced in 5 ms steps. Nevertheless, the influence of pacing history on the underlying cellular state may affect the CL at which discordant alternans develops and its spatial pattern.

Sex differences are implemented as ionic conductance scaling and did not explicitly include acute or chronic sex-hormone effects. SERCA was reduced uniformly to generate robust propagating alternans, representing a reduced Ca^2+^ handling reserve. Finally, the experimental optical mapping data presented only qualitatively support the physiological relevance of the predicted dissociation between CaT and APD phase. The central prediction that female strands maintain APD concordance across a CaT-discordant region more readily than male tissue remains to be tested experimentally in both sexes.

## CONCLUSION

Sex-dependent Ca²⁺-to-voltage coupling shapes alternans differently across spatial scales. Higher coupling gain in female myocytes amplifies cellular repolarization alternans, but in coupled strands, it strengthens electrotonic phase-locking and limits the translation of local Ca²⁺ discordance into APD discordance. This effect depends on electrotonic coupling and the spatial extent of β-adrenergic heterogeneity, supporting the hypothesis that spatial repolarization patterns emerge from competition between local Ca²⁺ instability and surrounding electrotonic influences. More broadly, these findings show that cellular electrophysiological phenotypes can be reshaped by multicellular interactions, underscoring the importance of sex, spatial heterogeneity, and intercellular coupling when translating cellular instability into arrhythmic risk.

## GENERATIVE AI STATEMENT

ChatGPT (OpenAI) was used during manuscript preparation to assist with language editing and refinement. Google Gemini and Claude (Anthropic) were used to support code verification and the conversion of code from MATLAB to C++. Consensus was used in addition to Google Scholar to identify potentially relevant literature on cardiac alternans. All AI-assisted outputs were critically reviewed, verified, and edited by the authors, who take full responsibility for the accuracy, integrity, and final presentation of the manuscript and figures.

## DATA AVAILABILITY

All source code will be made available on GitHub at https://github.com/drgrandilab.

## AUTHOR CONTRIBUTIONS

All authors conceived the work, drafted the work or revised it critically for important intellectual content, approved the final version to be published, and agree to be accountable for all aspects for the work.

